# Lessons from 20 years of plant genome sequencing: an unprecedented resource in need of more diverse representation

**DOI:** 10.1101/2021.05.31.446451

**Authors:** Rose A. Marks, Scott Hotaling, Paul B. Frandsen, Robert VanBuren

## Abstract

The field of plant genomics has grown rapidly in the past 20 years, leading to dramatic increases in both the quantity and quality of publicly available genomic resources. With this ever-expanding wealth of genomic data from an increasingly diverse set of taxa, unprecedented potential exists to better understand the genome biology and evolution of plants. Here, we provide a contemporary view of plant genomics, including analyses on the quality of existing plant genome assemblies, the taxonomic distribution of sequenced species, and how national participation has influenced the field’s development. We show that genome quality has increased dramatically in recent years, that substantial taxonomic gaps exist, and that the field has been dominated by affluent nations in the Global North and China, despite a wide geographic distribution of sequenced species. We identify multiple disconnects between the native range of focal species and the national affiliation of the researchers studying the plants, which we argue are rooted in colonialism--both past and present. However, falling sequencing costs paired with widening availability of analytical tools and an increasingly connected scientific community provide key opportunities to improve existing assemblies, fill sampling gaps, and, most importantly, empower a more global plant genomics community.

## Introduction

The pace and quality of plant genome sequencing has increased dramatically over the past 20 years. Since the genome assembly of *Arabidopsis thaliana*--the first for any plant--was published in 2000^1^, hundreds of plant genomes have been sequenced, assembled, and made publicly available on GenBank^2^ and other leading repositories for genomic data. With large, complex genomes and varying levels of ploidy, plant genomes have been historically difficult to assemble, but technological advances, such as long-read sequencing and new computational tools have made sequencing and assembly of virtually any species possible^3–5^. The number and quality of plant genome assemblies has increased exponentially as a result of these advances, enabling the exploration of both basic and applied research questions in unprecedented breadth and detail.

Land plants (Embryophyta) are extremely diverse and publicly available genome assemblies span over ~500 million years of evolution and divergence^6–8^. However, only a small fraction (~0.16%) of the ~350,000 extant land plants have had their genome sequenced, and these efforts have not been evenly distributed across clades^9^. For some plants (e.g., maize, *Arabidopsis*, and rice^10–12^) multiple high-quality genome assemblies are available, and thousands of accessions, cultivars, and ecotypes have been resequenced using high coverage Illumina data for these and other crop and model species^13^. Brassicaceae, a medium sized plant family, is the most heavily sequenced, with genome assemblies for dozens of species including *A. thaliana* and numerous cruciferous vegetables. In contrast, for most other groups, none or only a single species has a genome assembly. Ambitious efforts to fill taxonomic sampling gaps exist, including the Earth BioGenome and 10KP projects^14,15^ among others, but individual researchers can also play a role in expanding taxonomic representation in plant genomics.

The field of plant genomics is expanding rapidly, and a new generation of genomic scientists is being trained. Consequently, this is an ideal time to assess scientific progress while also developing strategies to increase equity and expand participation in the field. Economic disparities between nations, many of which were established due to colonialism, have a substantial impact on participation in science. Imperial colonialism provided scientists from the Global North access to a wealth of biodiversity, raw materials, and ideas that would have otherwise been inaccessible to them^16–18^. Over time, this led to a disproportionate accumulation of wealth and scientific resources in the Global North^19^, which contributed to the establishment and maintenance of global inequality^16,18,20^. Today, differences in funding, training opportunities, publication styles, and language requirements continue to drive similar patterns^18,21,22^. In plant genomics, the high costs of sequencing and provisioning computational resources are barriers to entry that perpetuate existing imbalances established due to colonialism and economic disparities. Luckily, the diminishing cost and increasing accessibility of sequencing technologies provides an opportunity to broaden participation and increase equity in plant genomics. This will require affluent nations and individuals to recognize their disproportionate access to biological and genetic resources and seek to increase participation rather than capitalizing on their economic privilege.

Here, we provide a high-level perspective on the first 20 years of plant genome sequencing. We describe the taxonomic distribution of sequencing efforts and build on previous estimates of genome availability and quality^23–26^. We show that an impressive and growing number of plant genome assemblies are now publicly available, that quality has greatly improved in concert with the rise of long-read sequencing, but that substantial taxonomic gaps exist. We also describe the geographic landscape of plant genomics, with an emphasis on representation. We highlight the need for the field, including its many affluent researchers and institutions, to work towards broadening participation. In our view, the wealth of publicly available plant genome assemblies can be leveraged to better understand plant biology while also continuing to decolonize a major field of research.

## Results

As of January 2021, 631 unique species of land plants had genome assemblies available in GenBank. We identified another 167 species with genome assemblies via literature searches and cross referencing against additional databases. If multiple genome assemblies were available for a species, we selected the highest quality genome assembly based on contiguity as a representative for that species. Unless otherwise noted, all analyses were conducted on the complete dataset of 798 genome assemblies (Table S1).

The number of plant genome assemblies has increased dramatically in the past 20 years, with marked improvements in quality associated with the advent of long-read sequencing (Fig. 1). Overall, 74% of plant genome assemblies were produced in the last 3 years. Contig N50 (the length of the shortest contig in the set of contigs containing at least 50% of the assembly length) also increased markedly in recent years, from 99.5 ±48.1 Kb in 2010 to 3,395.2 ±735.4 Kb in 2020. This increase appears to be driven primarily by advancements in sequencing technologies. Assemblies constructed with short-read technology (e.g., Illumina and Sanger) have significantly lower (p<0.0001) contig N50 (124.6 ±58.2 Kb) compared to those that incorporate long reads (e.g., PacBio and Oxford Nanopore) which have a contig N50 of 4,033.4 ±618.9 Kb. This difference translates to an impressive ~32x increase in mean contig N50 for long-read assemblies. Still, there are many extremely fragmented genome assemblies being published. Twenty-three of the genome assemblies in our dataset have a contig N50 below 1 Kb and 158 are below 10 Kb. These assemblies could be useful in some instances, but low-quality limits their value.

**Figure 1.**
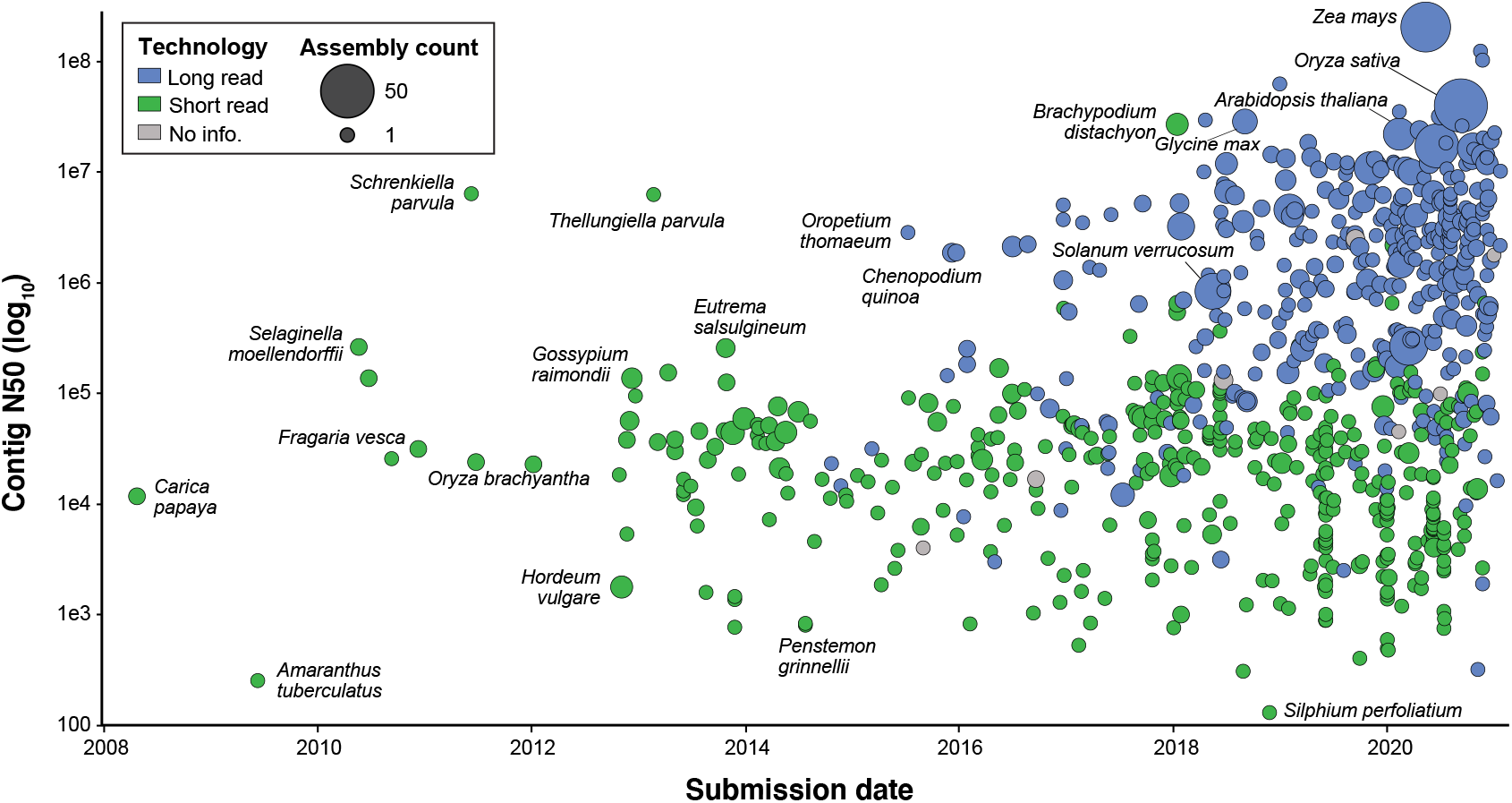
Changes in plant genome assembly quality and availability over time. Assembly contiguity by submission date for 798 plant species with publicly available genome assemblies. Points are colored by the type of sequencing technology used and scaled by the number of assemblies available for that species. There is an improvement in contiguity associated with the advent of long-read sequencing technology, and a noticeable increase in the number of genome assemblies generated annually. All assemblies generated prior to 2008 have since been updated and are therefore not included.

The first plants to have their genomes sequenced and assembled were model or crop species with simple genomes, but it is now feasible to assemble a genome for virtually any taxon. Still, taxonomic sampling gaps persist. Of the 137 land plant orders that have been described^27^, nearly half (76) lack a representative genome assembly. For the 62 orders with at least one genome assembly, a wide range of sampling depth is evident. For example, there are 83 species with genome assemblies in Brassicales, 80 in Poales, and 67 in Lamiales, yet there are 41 orders with fewer than 10 sequenced species. Six vascular plant orders are statistically overrepresented in genome assembly databases based on species richness. These include the agriculturally and economically important clades of Brassicales, Cucurbitales, Fagales, Malvales, Rosales, and Solanales. Four orders of vascular plants had significantly fewer genome assemblies than expected based on species richness (Fig. 2). Not surprisingly, these were speciose orders with significant ecological but comparatively less economic importance-- Asparagales, Asterales, and Gentianales—and the primarily polyploid order of Polypodiales (Fig. 2a). Bryophytes are poorly represented, with assemblies for only eight mosses, three liverworts, and three hornworts (Fig. 2a and Fig. S1). Diploid species are also statistically overrepresented in terms of genome assembly availability (Fig. 2b and Fig. S1) despite the widespread occurrence of polyploid plants^28^. Until recently, technological limitations have made it difficult to assemble high-quality polyploid genomes^4^. However, with the improvements that long-read sequencing technology offers, it is becoming more feasible to sequence and assemble complex polyploid genomes. As a result there are some highly contiguous tetraploid and reasonably contiguous hexaploid genome assemblies with mean contig N50’s of 1,855.7 ±474.3 Kb and 251.9 ±99.8 Kb respectively (Fig. S1).

**Figure 2.**
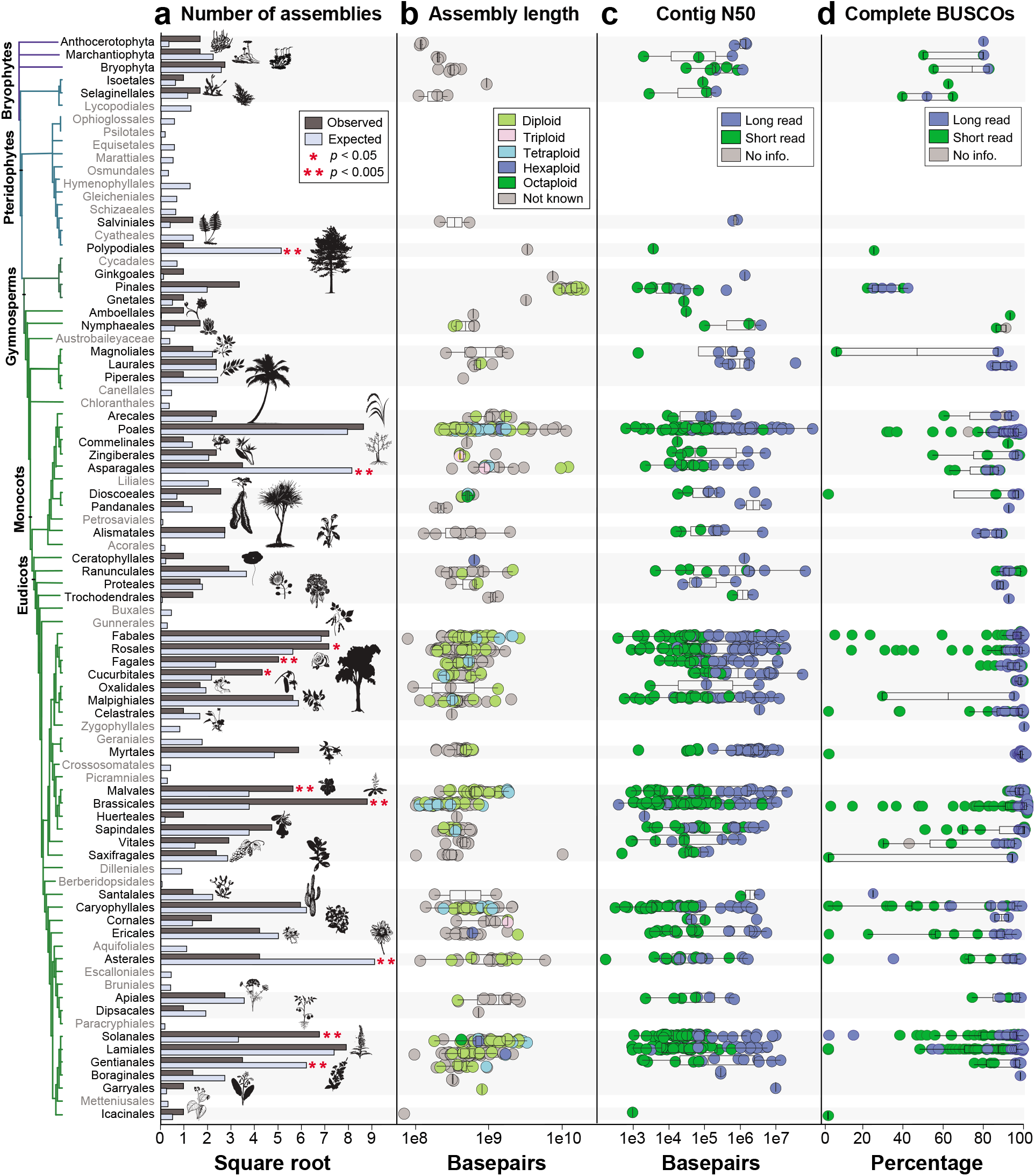
Comparison of genome availability and quality metrics for each land plant order. **a)** The number of species with publicly available genome assemblies as of January 2021 (n=798) versus the number expected for each order. Orders with no genome assemblies are shown in grey. Bryophytes are plotted at the phylum level but see Fig. S2 for bryophyte orders. Orders that showed a significant over- or under- representation are marked with asterisks. **b)** Length of assembly for each genome assembly. Points are colored by ploidy. **c)** Contig N50 for each genome assembly. **d)** Percentage of complete BUSCOs for each genome assembly. For (c) and (d), points are colored by sequencing technology.

To further assess differences in assembly quality and completeness, we quantified the percentage of Benchmarking Universal Single-Copy Orthologs (BUSCO; v.4.1.421) from the Embryophyta gene set in OrthoDB v.10^29^ that were present in each genome assembly deposited in GenBank. There was a high degree of variability in BUSCO completeness; percentages of complete BUSCOs (single and duplicated genes) ranged from 0 to 99% across the available genome assemblies (Fig. 2d). More contiguous genome assemblies with higher contig N50s had more complete BUSCOs (p<0.0088), and this correlation was associated with the use of long reads in the assembly process (p<0.0001; Fig. 2c-d and Fig. S3). Despite the wide range of assembly quality and completeness, no significant associations with genome size, taxonomy, or domestication status were identified.

We suspected that there has been a preference for sequencing economically important plants compared to wild or ecologically important species. To explore this, we classified the domestication status of each species with a genome assembly into six categories: (1) *domesticated*--plants that have undergone extensive artificial selection, (2) *cultivated*--plants that are used by humans but have not been subjected to substantial artificial selection, (3) *natural commodity*--plants that are harvested with little cultivation, (4) *feral*--plants that are not economically important but have still been influenced by human selection, (5) *wild*--plants that occur in the wild and have not been directly impacted by humans, and (6) *wild relatives*—wild plants that are closely related to domesticated or cultivated crops. Based on these categories, genome assemblies are available for 135 domesticated, 125 cultivated, 120 natural commodity, and 12 feral species. The remaining 406 genome assemblies are from wild species. Of these, 77 are wild relatives of crops (Fig. 3 and S4). While the number of human-linked species (domesticated, cultivated, natural commodity, and feral) with genome assemblies is largely equivalent to wild species, this equivalence reflects an extreme bias. There are far more wild (~350,000^30^) than domesticated species (~1,200-2,000 ^31,32^), suggesting that wild plants represent an untapped reservoir of genomic information.

**Figure 3.**
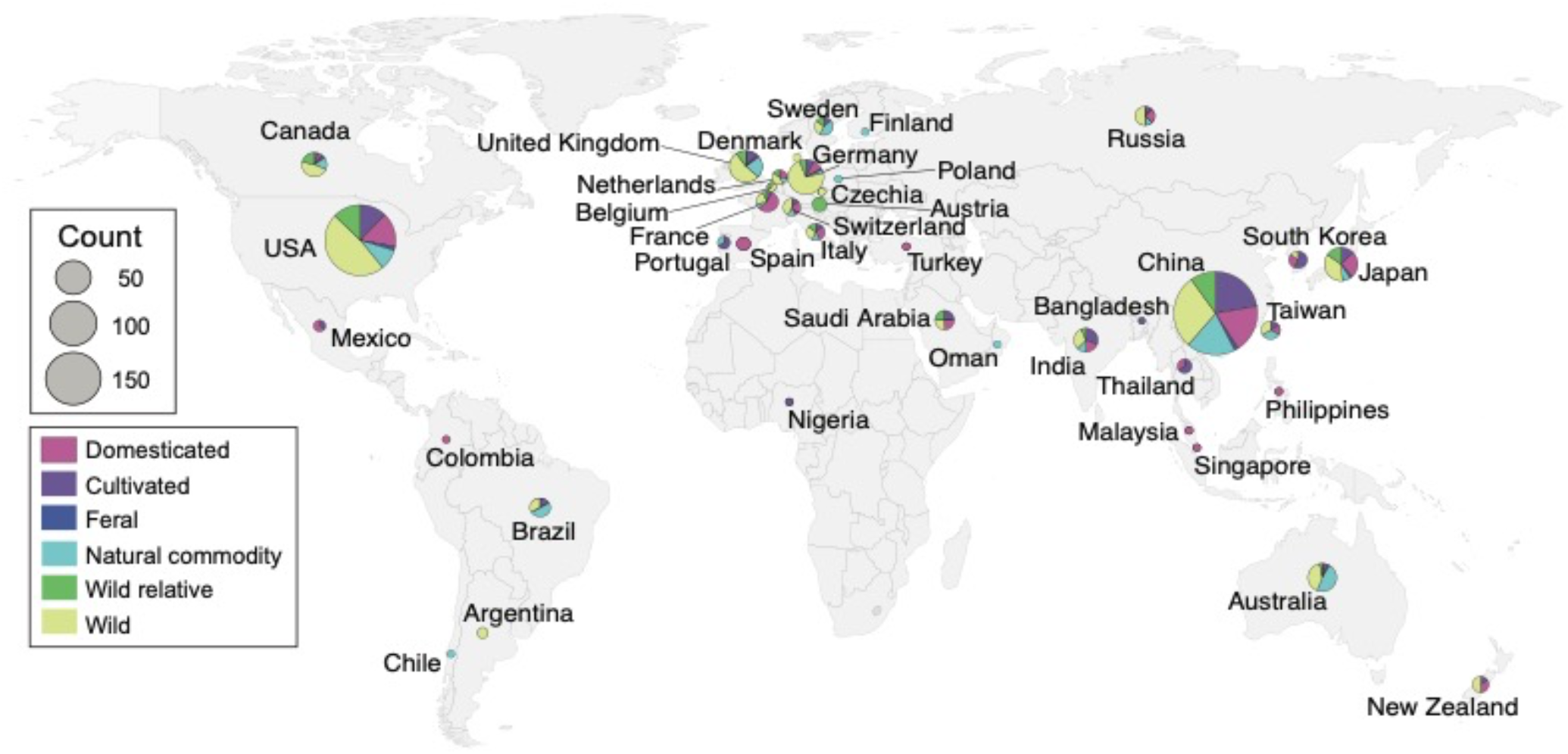
Geographic distribution of submitting institutions for plant genome assemblies. Circles are scaled by the number of genome assemblies produced in each nation and colored by the proportion of domesticated, cultivated, feral, natural commodity, wild, and wild relative species sequenced.

To better understand global participation in plant genomics, we identified the submitting institution for each genome assembly in our dataset. If the submitting institution was not listed, we identified the corresponding author for the associated publication and assigned the genome to the location of that institution. While this approach does not account for secondary affiliations in other nations, it does reveal where most of the scientific credit for a genome assembly is likely placed. We find that plant genome sequencing is dominated by China (233), the USA (212 assemblies), and Europe (165), with ~76% of genome assemblies attributed to one of those three regions (Fig. 3). Far fewer plant genome assemblies have been led by teams in Oceania (40), South America (9) and Africa (1). These patterns likely reflect well-documented differences in training incentives, facilities, and funding opportunities between nations^22,33–35^, many of which have been established and perpetuated through colonial practices^18^.

It is noteworthy that many of the sequenced plant genomes are of species that are native to or have economic importance in Africa and South America but have been sequenced elsewhere. We compared the center of diversity^36^ for all 135 domesticated crops in our dataset with the location of the institution that sequenced and assembled their genome. For this subset, we also investigated the affiliations of co-authors to gain insight into the extent of international collaborations. Although we did not account for geographical patterns of contemporary cultivation, the findings shed light on a disconnect between the origin of many crops and the institutions leading the genomic research on these species. There has been some reciprocal exchange between continents, but nearly all the crops native to Africa and South America have been sequenced off-continent. This represents a substantial global imbalance in genomics. There are dozens of major crops native to Africa and South America represented in GenBank, yet only one (*Phaseolus lunatus*) has a primary affiliation in South America and none were led by African institutions (Fig. 4). Even when co-author affiliations and collaborations are taken into account, this pattern holds true; most crops native to Africa and South America have been sequenced off-continent by non-collaborative teams. Specifically, most projects are led and conducted exclusively in the USA, China, and Europe. The lack of international collaboration is concerning, since it is likely that in some instances of off-continent genomics, sequenced material is chosen with minimal input from local stakeholders. Thus, the resulting genome assemblies may not represent the germplasm grown in production regions and the analyses my not address grower priorities. That being said, there are a growing number of inclusive and collaborative plant genomics projects such as the Orphan Crop Genome Consortium (http://africanorphancrops.org) and Africa BioGenome Project that are building capacity and broadening participation in plant genomics^22^.

**Figure 4.**
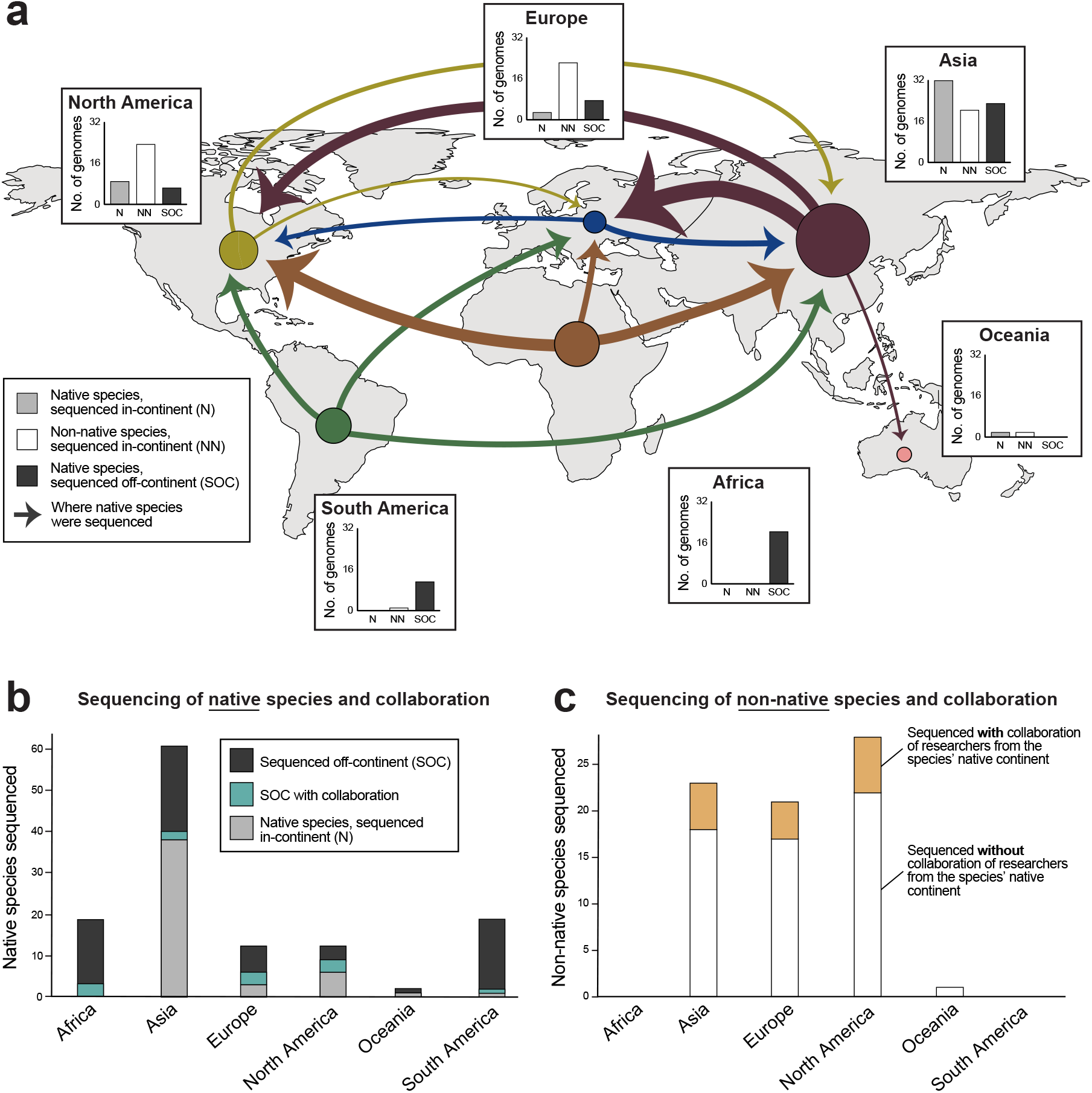
a) geographic perspective on where domesticated species (n=135) are native to versus where their genome assemblies were generated. Circle size and arrow weights are scaled by the number of genome assemblies being represented. The continents where arrows terminate represent where species were sequenced. **b)** The number of domesticated species native to each continent and the affiliations of the sequencing teams. **c)** The number of non-native species sequenced in each continent and the proportion of those efforts that included co-authors from the native range of the focal species.

## Discussion

The field of plant genomics has grown rapidly in the last 20 years, giving rise to an array of new tools, datasets, and biological insights. The quality of genome assemblies being produced today is much improved compared to even a few years ago, and this trend shows no signs of slowing. As has been observed for insects^37^, the improvement in plant genome quality appears to be driven largely by increased use of long-read sequences in assemblies. These technologies have enabled assembly of increasingly complex and polyploid genomes, opening up new arenas of research for plant genomocists. Despite these advances, major biases exist in both taxonomic sampling and participation. As the field continues to grow, there is an opportunity to fill key taxonomic gaps and build a broader, more representative discipline.

To date, plant genome scientists have focused mainly on sequencing economically important and model species with relatively simple genomes. This has led to major agricultural breakthroughs and fundamental scientific insights, and these densely sampled clades are ideal systems for investigating intraspecific variation and pan-genome structure. However, this approach has overlooked the wealth of information contained within the genomes of wild plants, which are extremely diverse, and largely untapped. Wild plants exhibit numerous diverse properties and produce a wide range of secondary compounds, many of which have important traditional and emerging pharmaceutical and industrial applications^38^. Numerous medical therapeutics and commercial materials are derived from or made to mimic plant-based compounds^39^, yet we have only begun to explore the rich chemical diversity of wild plants. Given the rapid loss of global biodiversity, it is critical that we take the opportunity to learn what we can from wild species before they disappear. Over the past ~100 years, we have witnessed a 60% increase in plant extinction^40^, and despite conservation efforts, this loss of biodiversity is projected to continue even under the most optimistic scenarios^41^. Improving genomic technologies provide an opportunity to explore, catalogue, and mine the immense diversity of information contained within wild species before they are lost.

In addition to taxonomic gaps, participation gaps are also prevalent in plant genomics. The field is dominated by a handful of affluent nations primarily from the Global North (e.g., USA, Germany, United Kingdom), and out analyses reveal a discrepancy between the native ranges of species and where their genomes are sequenced and assembled. In fact, 56% of all domesticated crops have had their genome sequenced outside of their continent of origin and only 13% included in-continent collaborators (Fig. 4). Much of the evolutionary innovation observed in landraces, locally adapted cultivars, and wild plants is exclusively maintained in the Global South, but only a handful of genome assemblies have been led by groups in those regions (with the exception of China, a notable economic and technological outlier relative to other nations of the Global South; Fig. 4). We argue that these dynamics are rooted in historical colonialism, economic barriers to entry, and are being perpetuated by contemporary “parachute science.” Historically, science was intimately linked to the rise of imperial colonialism^16–18^. Innovations in navigation and cartography enabled conquest of new territories by nations in the Global North and scientific curiosity actually motivated many early colonial expeditions^16^. Once colonies were established, they became the first sites for parachute science. Imperial scientists would travel to colonies, make collections, and take credit for “discoveries,” often discounting indigenous knowledge in the process. Over time, this led to a disproportionate accumulation of wealth, both scientific and economic, in the Global North that continues to drive disparities and participation imbalances today^18–20^. While historical colonialism set the stage for European nations to consolidate wealth and biological resources, both China and the USA have colonized surrounding territories in modern times. The resulting economic privilege has allowed these nations to capitalize on biological and genomic resources globally. Despite outward criticism of colonialism and legal provisions aimed at preventing international transport of biological and genetic resources (e.g., the Nagoya Protocol), affluent nations continue to lead bio- and genomic-prospecting efforts and parachute science remains prevalent^42,43^.

Going forward, we recommend that local communities and indigenous knowledge associated with the global reservoir of plant diversity^44,45^ form the backbone of plant genome collaborations. Currently, there are over a dozen plant genomics projects with African institutions as partners^22^, collaborative projects integrating indigenous knowledge^44^, and large-scale consortia with multinational participants are being established (e.g. the Africa BioGenome Project). These efforts all stand to broaden participation in plant genomics. As North American scientists, we acknowledge our own implicit and sometimes explicit participation in the sequencing and analysis of non-native plants. We encourage all plant scientists to strive to support local stakeholders, to incorporate indigenous knowledge into their work, and to invest in building systems and expertise for working with genomic resources in the location where they occur naturally. We believe that in-continent institutions should be encouraged to lead genome assembly of native species.

Plant genome science has arrived at an exciting moment with a rapidly expanding pool of genomic resources being generated by an increasingly diverse group of scientists. However, to take full advantage of the opportunities that a modern discipline affords and to ensure that the field continues striving for equity, we offer three recommendations. (1) Plant genome scientists should embrace long-read sequencing technologies and leverage them whenever possible to generate new assemblies. This is already occurring but given the massive disparity in quality between assemblies generated with short-read versus long-read data, the need for continued adoption cannot be overstated. (2) Despite considerable progress, the taxonomic scope and domestication status of plants with available genome assemblies should continue to be expanded. In our view, attention should be focused on generating assemblies for clades that have none (e.g., Hymenophyllales, Cyatheales, Geraniales, Dilleniales; see Fig. 2a), adding more complex plant genome assemblies (e.g., repetitive and/or polyploid), and sequencing wild species. (3) While the progress driven by large-scale consortia is undeniable, it is important that researchers in the discipline are mindful of the signatures of colonialism--past and present--in plant genome science. To this end, we should collectively monitor consortia, collaborations, and projects to ensure that ethical approaches are being taken, in-country peoples are given a voice, and that participation and access to resources is broadened at every level. Ultimately, a diverse, thriving discipline with empowered researchers across continents regardless of socioeconomic status will yield the greatest potential to meet the economic, social, and evolutionary challenges facing 21st century plant science.

## Methods

Species and assembly metadata are provided in Table S1. To compile the best genome assemblies for all land plants we downloaded the most contiguous genome assembly for each species represented in GenBank, in January 2021. Genome assemblies were downloaded using the *download-genome* function of NCBI’s datasets tool (v.10.9.0), and metadata were extracted using the *assembly-descriptors* function of NCBI’s datasets tool. Data on sequencing technology, coverage, assembler, and submitting institution were retrieved using the python script *scrape*_*assembly*_*info.py* (https://github.com/pbfrandsen/insect_genome_assemblies). For genome assemblies with no reported sequencing technology on GenBank, we went to the publication associated with the assembly (if available) and identified the sequencing technology from the reported methods. In addition, we conducted an extensive literature search to identify additional genome assemblies not deposited in GenBank. To do so, we took advantage of review papers summarizing plant genome assemblies^23–26^ and other datasets such as PlaBi database (www.plabipd.de), Phytozome (https://phytozome.jgi.doe.gov/), Fernbase (https://www.fernbase.org/), and (https://en.wikipedia.org/wiki/List_of_sequenced_plant_genomes). We cross referenced these datasets against NCBI to develop a nonredundant but comprehensive list of plant genome assemblies. For these genome assemblies not deposited in NCBI, metadata (including assembly size, contig N50, sequencing technology, authorship, and domestication status) was extracted from the primary publication.

Higher level taxonomy for each species was integrated with taxonkit^46^. To place species in a phylogenetic context, we identified the most up-to-date phylogenies for each major group of land plants and grafted them together. For angiosperms we used the APG IV tree^47^, for gymnosperms and pteridophytes we used the APGweb tree (http://www.mobot.org/MOBOT/research/APweb), and for bryophytes we used iTol^48^. Many of the relationships among these groups are still poorly resolved or under ongoing revision, but for the purposes of this work, they are sufficient to visualize general relationships among clades.

To identify cases where the observed number of genome assemblies for an order differed significantly from the expected number based on species richness, we tested for an over- or under-representation of genome assemblies in each land plant order using Fisher’s Exact Tests. To do so, we compiled a list of the total numbers of species in each land plant order. For vascular plants, we used the Leipzig Catalogue of Vascular Plants (LCVP; v 1.0.3)^27^ in combination with the summaries provided in^49^. For bryophytes, we compiled data from the Plant List (http://www.theplantlist.org; accepted names only) and cross referenced these against the Missouri Botanical Gardens *Index of Bryophytes* (http://www.mobot.org/mobot/tropicos/most/bryolist.shtml). Next, we computed the number of genome assemblies that would be expected for each order if sampling effort was evenly distributed. We then ran Fisher’s Exact Tests to identify clades with a statistical over- or under-representation of genome assemblies.

To quantify the distribution of polyploid genome assemblies, we pulled data on chromosome number and ploidy from the Kew Botanical Garden’s plant C-values database^50^. These data were used to calculate the total reported number of species with each ploidy level. We then calculated the number of genomes assemblies expected for every ploidy level. Using these numbers, we ran Fisher’s Exact Tests to identify ploidy levels that had an over- or under-representation of genome assemblies.

We classified the domestication status of each species using a six-category scale. Each species was designated as either (1) *domesticated*--plants that have undergone extensive artificial selection, (2) *cultivated*--plants that are used by humans but have not been subjected to substantial artificial selection, (3) *natural commodity*--plants that are naturally harvested with little cultivation, (4) *feral*--plants that are not economically important but have still been influenced by human selection, (5) *wild*--plants that occur in the wild and have not been directly impacted by humans, and (6) *wild relatives*--plants that are closely related to domesticated or cultivated crops. Using this classification system, we computed the total number of genome assemblies for each category.

We investigated the completeness of genome assemblies by quantifying the percentage of complete, fragmented, and missing Benchmarking Universal Single-Copy Orthologs (BUSCO; v.4.1.421) from the Embryophyta gene set in OrthoDB v.10^29^. We ran BUSCO (v.4.1.4) in --*genome mode* on each GenBank assembly with the --*long* option. We did not include the genome assemblies gathered from published papers in these analyses due to difficulties in accessing the genome files. We tested for an association between the percentage of complete BUSCOs (single and duplicated) and the contiguity of genome assemblies (contig N50) using a linear model. Similarly, we tested for an effect of sequencing technology on the percentage of complete BUSCOs with the assembled length size included as a random effect.

To estimate the geographic distribution of plant genome projects, we identified the submitting institution for each genome assembly in our dataset. If the submitting institution was not listed, we identified the corresponding author for the publication and assigned the genome to the location of that institution. Next, we compiled data on the center of diversity^36^ for all 135 domesticated crops with genome assemblies. For these species we dissected authorship in more detail, in order to account for collaborative efforts. We looked at the affiliations of all authors for each publication relative to the center of diversity of the sequenced species. Projects were scored as either *“in-continent team”*, *“off-continent team”*, *“led by off-continent team*, *with in-continent contributions”*, or *“led by in-continent team*, *with off-continent contributions”*. Using these categories, we summarized global patterns of plant genome sequencing relative to the center of origin for these important crops.

## Acknowledgements

This work was supported by an NSF Postdoctoral Research Fellowship in Biology (PRFB-1906094) to RAM and NSF grant MCB-1817347 to RV. SH was supported by NSF award OPP-1906015. The Plant Resiliency Institute at Michigan State University provided additional funding that supported this work.

