## Supplementary information for "Lessons from 20 years of plant genome sequencing: an unprecedented resource in need of more diverse representation" (from another version of this preprint: version 1, 2021-05-31)

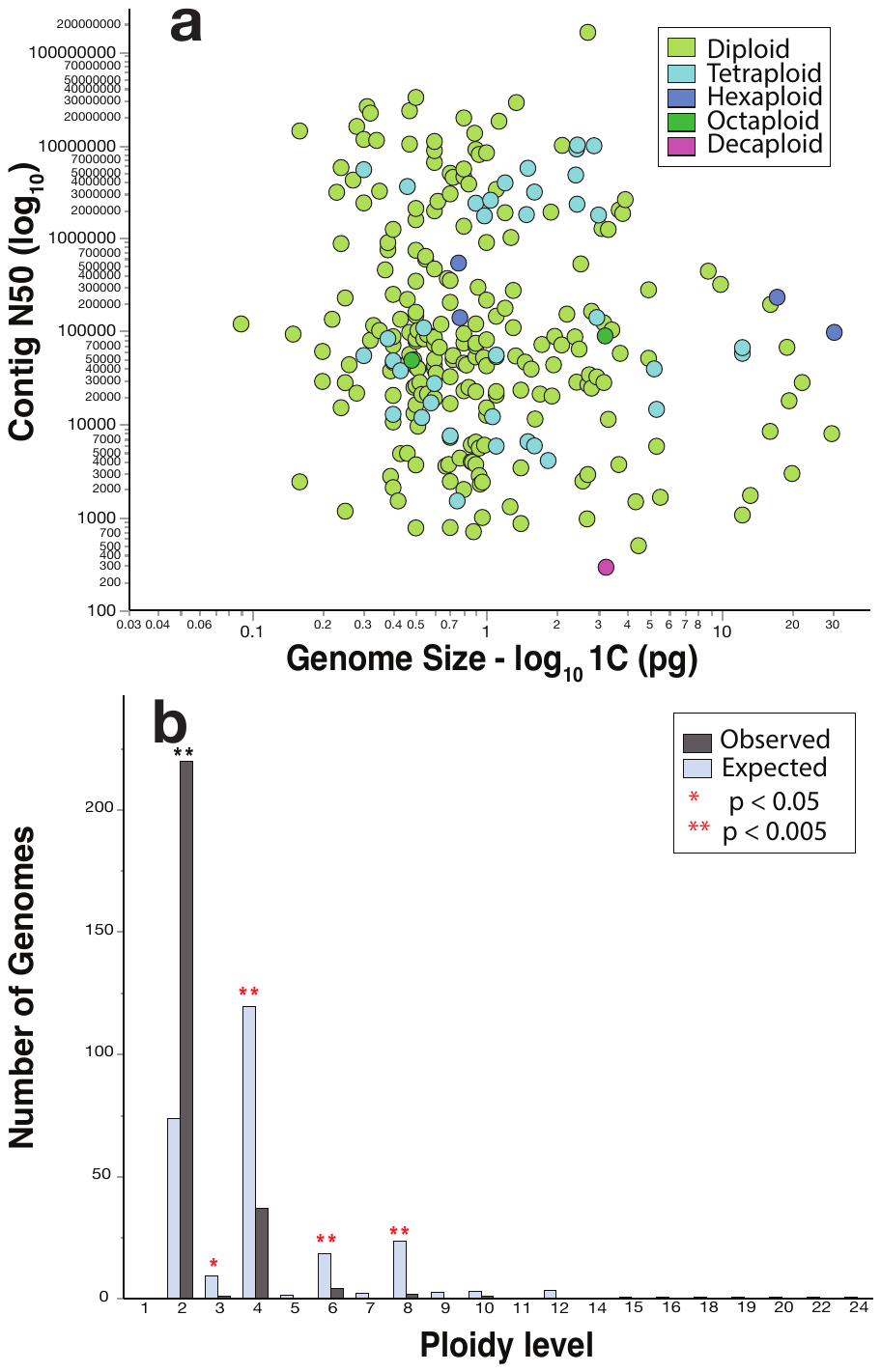

**Figure S1. a)** The genome contiguity (N50) by estimated genome side, with the ploidy of each species indicated by color. Contiguity is not associated with differences in genome size and polyploid species are reasonably well assembled. **b)** The observed vs expected number of genome assemblies available for each ploidy level. Orders that showed a significant over- or under-representation are marked with ** (p < 0.005) or * (p < 0.5).

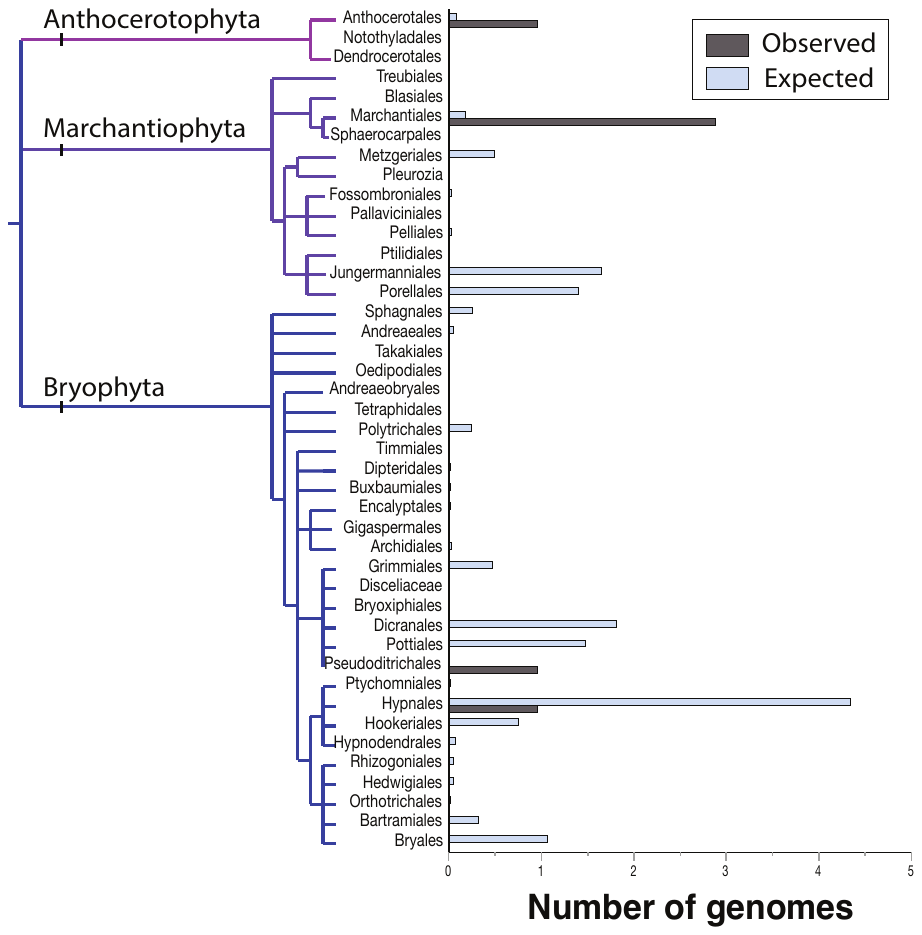

**Figure S2.** The number of species with publicly available genome assemblies versus the number expected for each bryophyte order. Orders with no genome assembly are in grey. Orders that showed a significant over- or under-representation are marked with ** (p < 0.005) or * (p < 0.5)

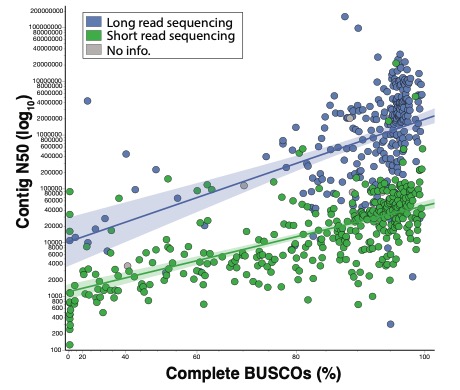

**Figure S3.** Genome contiguity is positively associated with the percent of complete BUSCOs identified. Overall, assemblies generated with long read sequencing capture a higher percentage of complete BUSCOs.

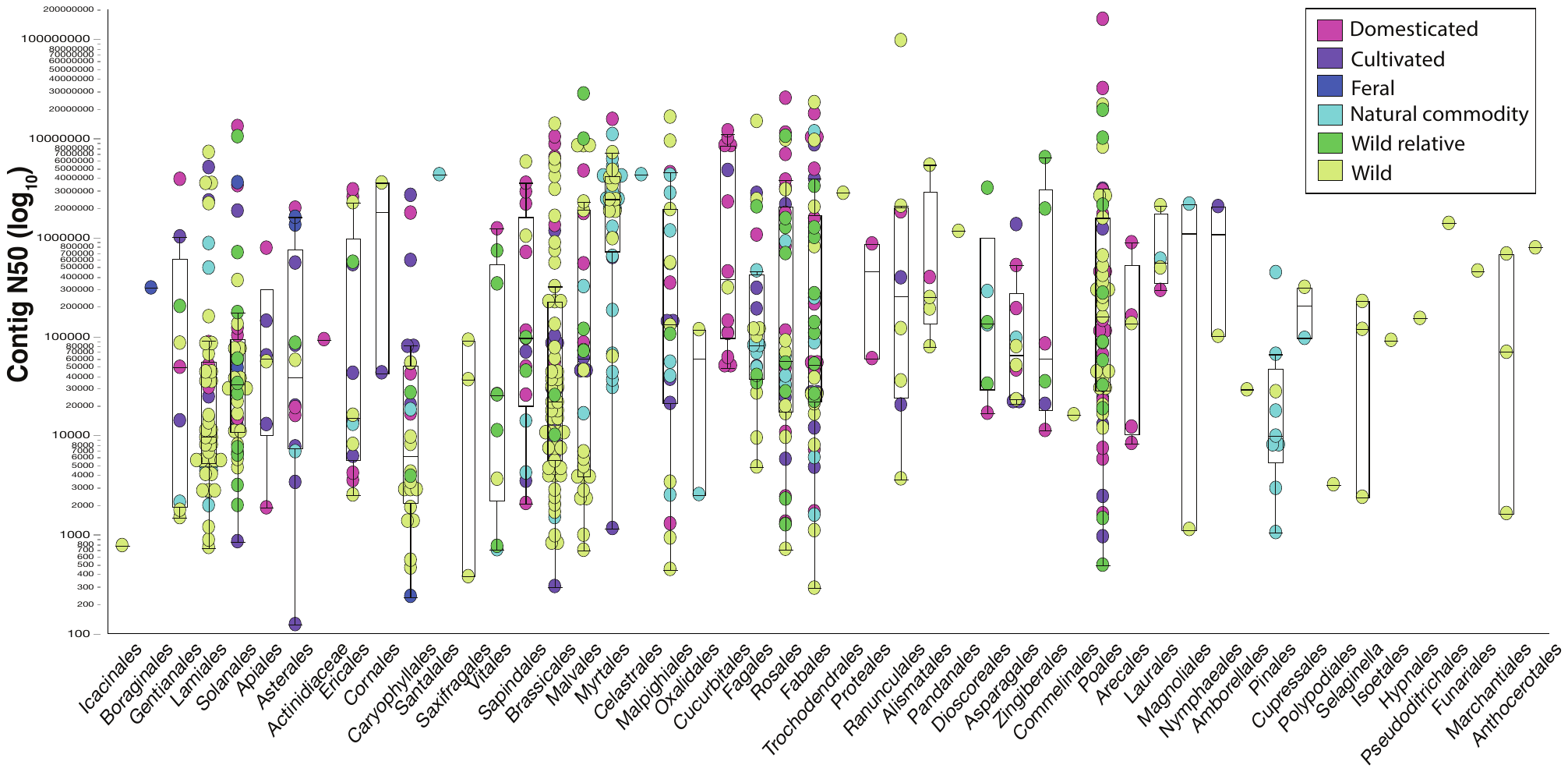

**Figure S4.** The distribution of sequenced domesticated, cultivated, natural commodity, feral, wild, and wild relative plants across plant orders and their contiguity.
